# Impaired temporal prediction mechanisms in dyslexia

**DOI:** 10.64898/2026.01.16.699956

**Authors:** Pierre Bonnet, Barbara Tillmann, Eden Chettih, Nathalie Bedoin, Anne Kösem

## Abstract

Effective speech analysis involves deconstructing the acoustic signal into identifiable linguistic units, which depends on the ability to recognize and anticipate temporal patterns within the speech stream. However, these processes may be less efficient in individuals with dyslexia. This study investigated the effects of temporal context and related temporal predictions in dyslexic adult participants and matched control participants, using an auditory oddball task with non-verbal stimuli. Pure tones were presented in sequences, and participants were requested to discriminate the pitch of target stimuli. The temporal intervals between the sounds varied in regularity across the sequences, thereby creating contexts with different levels of temporal predictability. At the end of each sequence, participants were prompted to evaluate the perceived rhythmicity of the sequence and to assess their own performance in the auditory discrimination task. Dyslexic participants demonstrated overall lower accuracy in discriminating target sounds than controls. They also showed reduced influence of the temporal context of the sequences on response times, while controls responded faster in sequences that were temporally more regular and predictable. Additionally, individuals with dyslexia perceived the rhythmicity of sound sequences less accurately, overestimating the temporal regularity in irregular sequences and underestimating it in regular sequences. They also reported lower overall confidence in their ability to perform the task compared to control participants. Altogether, these findings provide converging evidence for altered temporal prediction abilities in dyslexia, which may impact auditory perception and then impair language processing.

## INTRODUCTION

Dyslexia is a neurodevelopmental disorder that affects an individual’s capacity to fluently read, write, and spell. Research suggests that dyslexia is unlikely to result from a single cause; rather, it may stem from a combination of multiple risk factors (Pennington, 2006; van Bergen et al., 2014). Reading acquisition indirectly depends on the ability to segment spoken language into a sequence of perceptually discrete linguistic units to construct a reliable phonological system. Impairments in temporal auditory segmentation processes may therefore contribute to reading difficulties associated with dyslexia (Goswami, 2011; Lense et al., 2021). Consistent with this interpretation, dyslexia is often characterized by phonological processing deficits (Cole et al., 2020; Shaywitz & Shaywitz, 2005), as well as difficulties in processing auditory information over time (Farmer & Klein, 1995; Habib, 2021; Meilleur et al., 2020; Tallal, 2004). Children and adults with dyslexia tend to demonstrate lower performance in temporal processing tasks, whether these tasks focus explicitly on timing (e.g. judgments on duration/temporal order and simultaneity (Casini et al., 2017; Tallal et al., 1993; Tallal & Piercy, 1973), rhythm discrimination (Bégel et al., 2022; Fiveash et al., 2021; Overy et al., 2003; Ozernov-Palchik et al., 2018)) or whether they rely on implicit temporal cues (such as phoneme discrimination or production based on durational contrast (Bouhon et al., 2023; Serniclaes et al., 2021; Casini et al., 2017), or electrophysiological mismatch negativity response to duration-deviant non-verbal and verbal sounds (Corbera et al., 2006; Huttunen-Scott et al., 2008; Stefanics et al., 2011)). Furthermore, temporal task abilities of infants and children can predict language skills later in development (Benasich & Tallal, 2002; Bonacina et al., 2021; Kertész, & Honbolygó, 2023; Nitin et al., 2023; Ozernov-Palchik et al., 2018; Plourde et al., 2017; Steinbrink et al., 2014).

Our present study tested whether the altered temporal processing in dyslexia could stem from an impairment in temporal prediction mechanisms (Fiveash et al., 2021; Ladányi et al., 2020; Taha et al., 2025). Music and speech contain temporal regularities (Ding et al., 2017; Fiveash et al., 2021) that could induce temporal expectations about when the next item will occur. Temporal expectations are known to facilitate the processing of sensory stimuli that align with rhythmic regularity (Jones, 2019; Large & Jones, 1999; Nobre et al., 2012). This processing facilitation is thought to arise from the synchronization between sensory temporal regularities and low-frequency neural oscillations (Cravo et al., 2013; Herbst & Obleser, 2017; Miniussi et al., 1999; Schroeder & Lakatos, 2009; Wilsch et al., 2020). Neural oscillations in the delta (1–4 Hz) and theta (4–8 Hz) ranges have been identified as contributing to the temporal processing of speech (Giraud & Poeppel, 2012; Kösem & van Wassenhove, 2017; Zoefel & Kösem 2024). They have been shown to reflect temporal expectations based on syllabic and phrasal rates, which influence how the continuous speech signal is parsed and understood (Bree et al., 2021; Guiraud et al., 2018; Kösem et al., 2018; 2020). Therefore, inefficient temporal processing in dyslexia has been hypothesized to originate from impaired neurophysiological tracking mechanisms carried by neural oscillations (Chang et al., 2021; Goswami, 2011, 2018; Lallier et al., 2017). Dyslexia would be associated with atypical speech-brain synchronization, which would lead to challenges in accurately segmenting syllables and words in the continuous speech signal and analyzing its prosodic contour information. These auditory processing limitations would in turn significantly affect reading proficiency (Lallier et al., 2017; Goswami, 2022). In line with this hypothesis, several reports have shown that individuals with dyslexia exhibit reduced or altered synchronization of neural oscillations to temporally regular sequences within the delta and theta frequency ranges when compared to neurotypical participants (Destoky et al., 2022; Fiveash et al., 2020; Guiraud et al., 2018; Hämäläinen et al., 2012; Leong & Goswami, 2014, 2014; Lizarazu et al., 2021; Molinaro et al., 2016; Soltész et al., 2013; Thomson et al., 2006; Thomson & Goswami, 2008). Yet, the relationship between reduced speech-brain synchronization and deficits in temporal prediction remains indirect and definitive evidence for impairments in temporal prediction mechanisms per se is lacking in individuals with dyslexia (Kösem, Dai, et al., 2023; Zoefel & Kösem, 2024). Recent behavioral studies suggest that dyslexia may be associated with difficulties in the temporal prediction of auditory information (Pagliarini et al., 2020). However, these findings have primarily been investigated through tapping tasks where temporal information is processed explicitly, and further research is needed to determine the extent to which temporal predictions implicitly impact auditory perception in dyslexia.

Beyond the temporal regularities in speech, its acoustical structure also encompasses a range of temporal variations (Cummins, 2012; Jadoul et al., 2016; Singh & Theunissen, 2003; Varnet et al., 2017). Gaining insight into how temporal prediction mechanisms operate in the presence of irregularities is therefore essential for understanding their impact on continuous speech processing. This was the aim of a recent behavioral study that revealed that typical listeners can develop temporal predictions also in auditory contexts that are not fully rhythmic and less predictable (Bonnet et al., 2024). Participants were asked to discriminate between standard and deviant sounds that were embedded in 3.5 min-long sound sequences. The Stimulus Onset Asynchronies (SOAs) between the sounds of the sequences were drawn from distributions with the same mean (500 ms) but with distinct temporal variability (Fig. 1). Results showed that the statistical properties of non-periodic auditory sequences influenced auditory discrimination performance. Specifically, participants’ discrimination accuracy was enhanced and response times were faster in sequences with higher temporal regularity (periodic contexts) compared to more variable temporal sequences. The findings indicate that temporal prediction mechanisms are effective in probabilistic contexts, i.e. when sound sequences are not strictly periodic. Additionally, the influence of temporal statistics tends to diminish progressively as the temporal variability of the sound sequence increases. Our present study uses this paradigm, originally designed for typically developing adults, to investigate temporal prediction abilities in individuals with dyslexia. Based on previous findings suggesting altered/impaired temporal prediction mechanisms in dyslexia, performance of dyslexic participants was expected to show a reduced benefit from the temporal regularities present in the context relative to control participants.

**Figure 1:**
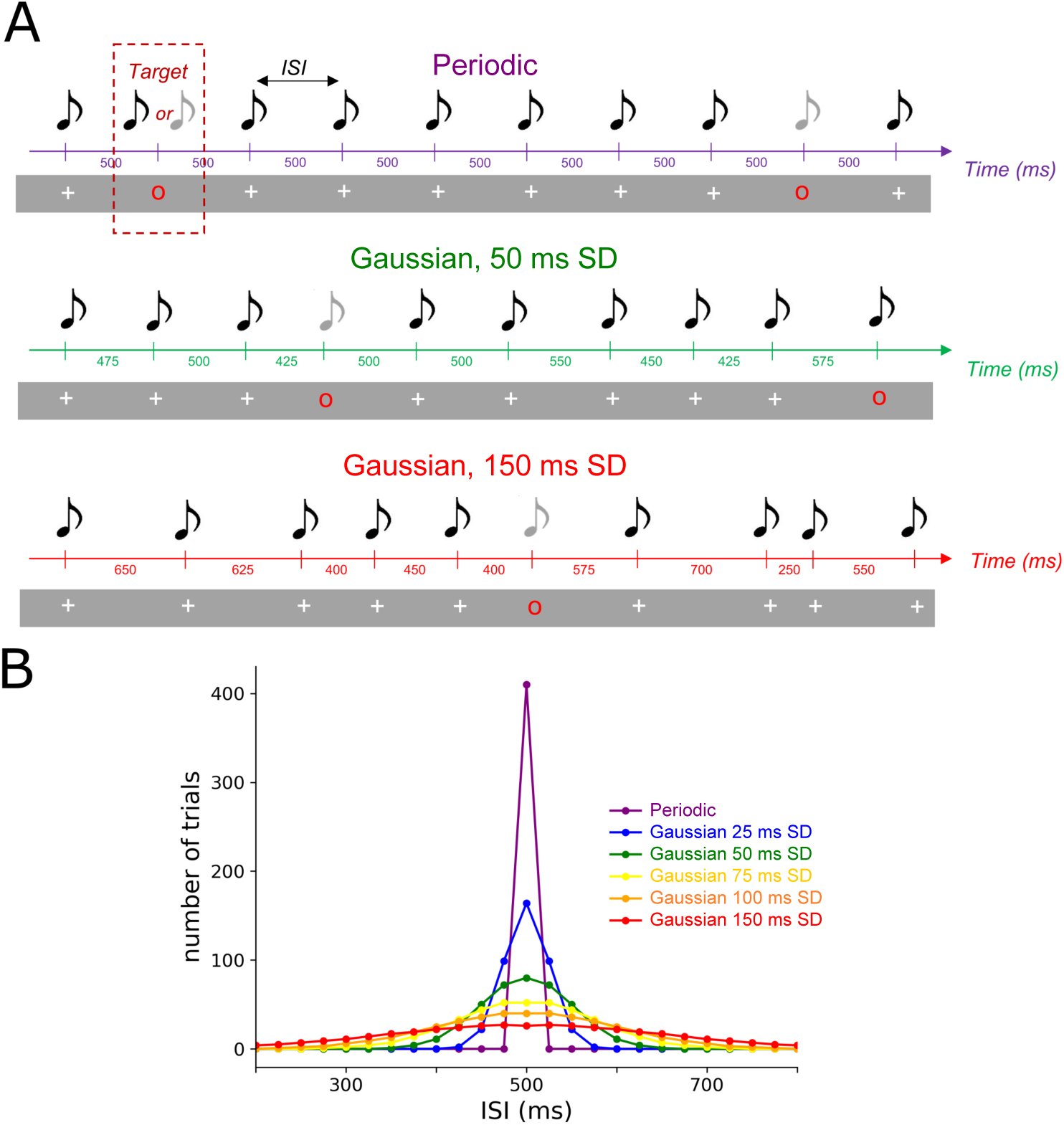
**Experimental design**. (A) Example of three sequences of different temporal standard deviations (SD) used in this experiment, adapted from (Bonnet et al., 2024). Each sequence consisted of a stream of simultaneous auditory and visual stimuli. A white cross indicated the simultaneous presentation of a standard auditory stimulus (440 Hz pure tone). A red circle indicated the presentation of a target auditory stimulus, on which participants had to discriminate between a standard (440 Hz) and deviant (220 Hz) pure tone. (B) Distribution of SOAs in each sequence. For each sequence, the distributions of the SOAs were drawn of Gaussian distributions with equal means (SOA = 500 ms) but distinct SDs. Six conditions were designed: from 0 (periodic) to 150 ms SD.

## METHODS

### Participants

Twenty-seven dyslexic adults (19 women, mean age = 24.33 years ± 6.29 *SD*) and twenty-six control participants (19 women, *M* = 24.14 years ± 6.29 *SD*) took part in the experiment. Dyslexic participants were required to have received their diagnosis from a speech therapist for at least two years during childhood. The persistence of dyslexic traits for each participant was further validated with French evaluation tools designed to assess language skills in individuals aged over 16 years (see Procedure and Results section). All participants were screened to exclude any hearing, neurological or psychiatric disorders. Participants reported not having taken private music or dance lessons, either currently or in the past. Furthermore, they were included only when they self-reported the absence of other neurodevelopmental disorders commonly associated with dyslexia, including other DYS-disorders such as dyscalculia, dysphasia, and dyspraxia, as well as attention deficit disorder with or without hyperactivity (ADHD), for which the participants also completed an ADHD symptom screening questionnaire (ASRS-v1.1) (Kessler et al., 2005), as this comorbidity may impact task performance independently of the dyslexic disorder (Calderone et al., 2014; Gabrieli, 2009). One dyslexic participant was excluded because of an ADHD score exceeding the established test criteria (Part A score > 4, Part A contains the six questions most predictive of ADHD (Kessler et al., 2005)). Thus, 52 participants were included in the study. The study was approved by a national ethics committee (CPP 220 B17), and all participants signed informed consent before starting the experiment.

### Stimuli

Stimuli of Bonnet et al. (2024) were used. The stimuli consisted of standard tones (pure tones at 440 Hz with 5 ms ramp onset/offset) and deviant tones (pure tones at 220 Hz with 5 ms ramp onset/offset) (see Figure 1). Visual cues were presented simultaneously to each sound, they appeared at the onset of the sound and disappeared at their offset. When a white cross was displayed, the synchronized auditory stimulus was always a standard sound. When a red circle appeared, participants were required to respond to which sound they heard by pressing a key on the keyboard (left key for a standard sound and right key for a deviant sound). The inter-stimulus intervals (SOAs) between the sounds of the sequence were drawn from distinct Gaussian distributions of the same mean (500 ms) but of distinct standard deviation (SD). Six types of sequences with different SOA distributions were presented: periodic condition (with 0 ms SD), and gaussian distributions with 25 ms, 50 ms, 75 ms, 100 ms, and 150 ms SD. Each sequence comprised 410 tones, with 56 of them synchronized with the red circles (targets). Between two targets, a minimum of four and a maximum of ten standard tones were presented. Sound sequences were delivered alongside continuous white noise. The signal-to-noise ratio between the sound sequences and the white noise was modified using a staircase procedure (see Procedure section). All stimuli were generated and presented using Psychopy software version 3.0.2 (Peirce et al., 2019).

### Procedure

Prior to the main experiment, dyslexic and control participants completed various questionnaires to assess dyslexia, ADHD and IQ.

Reading skills in adults were assessed with four sub-tests of the ECLA 16+ battery (Gola-Asmussen et al., 2010), a tool designed to screen adults for dyslexia in French. The tests took approximatively 10 minutes.

I. *Reading test “L’Alouette”.* This read-aloud test indicates reading speed and errors without possible anticipation of semantic information. The results obtained were compared with the mean, SD and percentiles according to the participant’s age (Lefavrais, 1967, 2005; Cavalli et al., 2018).
II. *Word dictation*. A dictation test with 10 orthographically regular words, 10 irregular words and 10 pseudowords was used to investigate participant’s lexical and phonological writing difficulties.
III. *Initial phoneme deletion test*. This test of phonological awareness analyzes the ability to extract syllables and phonemes, and to manipulate them deliberately. Participants were asked to remove the first sound of a specific word (e.g. “cliché” > “liché”)
IV. *Spoonerism test*. This phonological memory test requires participants to isolate and swap the first phoneme of two words, presented in pairs (e.g. “dossier”-“massage”), and to produce the resulting words (e.g. “mossier”-”dassage”).

To assess IQ, participants completed a simplified version of the Raven Matrices test (Bilker et al., 2012) (9-item version A11, B12, C4, C12, D7, D12, E1, E5, E7). The Raven Matrices are a multiple-choice non-verbal intelligence test consisting of matrices to be completed with progressive difficulty (QI SPM; Standard Progressives Matrices - Raven, 1936). The test took approximatively 5 minutes to be completed.

In addition, each participant completed an ADHD symptom screening questionnaire (ASRS-v1.1) (Kessler et al., 2005) comprising 18 questions aligned with the DSM-IV-TR criteria and providing a list of symptoms that may indicate the presence of ADHD. The 18 items are divided into two parts: Part A (6 questions), and Part B (12 questions). Part A of the questionnaire is the fundamental component for screening with the ASRS v1.1, while Part B offers additional insights to guide therapeutic follow-up. We restricted our analyses to Part A because it is the most indicative for making a diagnosis of ADHD and constitutes the best tool for screening this disorder. Data from participants who scored above the established exclusion criterion from the test (score superior to 4 in Part A) were excluded (1 dyslexic participant). The test took approximately 5 minutes to be completed.

After having completed these tests, participants were seated in a soundproof experimental room approximately 70 cm from a computer screen and wore headphones. The experiment began with a staircase adjustment procedure lasting about five minutes: the staircase procedure aimed to ensure that participants understood the instructions and to adjust the sound intensity to an optimal level. By adjusting the loudness of the stimuli and modifying the signal-to-noise ratio between the sound sequences and the white noise background, we established a condition that resulted in an average performance of approximately 80% correct pitch discrimination responses of targets. In the staircase, 75 target sounds were presented in temporally irregular sequences with SOAs drawn from a gaussian distribution with 500 ms mean and 150 ms SD, and target trials (sounds associated with a red circle visual cue) could occur every ∼3-6 tones. During the staircase procedure, the script paused after each target presentation to await the participant’s response before proceeding with the sound sequence. The resulting mean SNR across participants was of −13.87 dB, within [−15.89, −9.58] dB range, comparable with our previous study (Bonnet et al., 2024).

Once the staircase had been completed, the experiment started, with the same experimental procedure as in Bonnet et al. (2024). The experimental phase contained 12 blocks, each block lasting 3.5 min. Each block consisted of a sequence of auditory tones masked in constant noise, and the tones were associated with visual cues (a white cross or a red circle). For each block, when a red circle was presented on the screen (target trials), participants were asked to judge whether the sound heard through the headphones was of the same pitch than the tones presented with the white cross (standard sound) or whether it was different in pitch (deviant sound), by pressing two different keys on the keyboard (Fig. 1A). To vary the temporal regularities of the context, the SOAs between the sounds of each block was drawn from distinct distributions. In the Periodic condition, the SOA was fixed at 500 ms. For the Gaussian conditions, the SOAs were drawn from Gaussian distributions with distinct SDs of 25 ms, 50 ms, 75 ms, 100 ms and 150 ms (Fig. 1B). Each block consisted of 410 tones including 56 target trials. Between two target trials, a minimum of four standard sounds and a maximum of 10 standard sounds (uniform distribution) could occur. The target trials could not appear in the first 10 sounds of the sequence. Therefore, 112 target trials were obtained for each condition. Between each block, participants could take breaks to maintain optimal alertness and concentration throughout the experiment. Two blocks per condition were presented to participants in a pseudo-randomized order to prevent the same condition from occurring twice in succession. After completing each block, participants were asked to subjectively evaluate (1) their perception of the overall temporal regularity of the sound sequence they had just heard, using a scale from 0 (completely irregular) to 10 (completely periodic) (i.e. “ do you think that the sounds in the sequence are presented at a regular pace? from zero = not regular at all, to 10 = completely periodic)”, and (2) their own performance in that block, rated on a scale from 0 (random responses) to 10 (completely accurate/perfect).

### Statistical analyses

Statistical analysis of the data was conducted using R Studio (R 4.1.2 (2021-11-01)). Student’s *t*-tests and chi-square (χ^2^) tests were performed to compare the demographics of the two groups. For the auditory discrimination task, we performed statistics on deviant discrimination accuracy and response times (RTs). We also analyzed the subjective rhythmicity and subjective performance ratings given at the end of each auditory sequence. One dyslexic participant was excluded for the rating analyses due to a technical issue during data acquisition. Generalized linear mixed models (GLMMs) using lme4 (version 1.1-28) (Bates et al., 2014) were employed on four dependent variables: accuracy (1 for a correct response, 0 if incorrect, binomial distribution), log-transformed RTs (gamma distribution), subjective temporal regularity rating (normal distribution), and subjective performance rating (normal distribution). Trials with RTs shorter than 0.1 s and longer than 3 s were excluded from the analysis. For accuracy, rating on rhythmicity and rating on performance, we performed a model with the fixed effects of the Group condition (dyslexic, control), the context’s Temporal Variability (continuous variable ranging from 0 ms to 150 ms Temporal *SD*), their interaction, and Participant was included as a random factor. For RTs, we additionally incorporated the response Correctness (1 for correct, 0 for incorrect) and their interaction with Group and Temporal variability as factors in the model.

Model*_Accuracy_* ∼ Group * Temporal Variability + (1|Participant)
Model*_RT_* ∼ Group * Temporal Variability * Response Correctness + (1|Participant)

To test the impact of ADHD score on performance, we compared the base models to models including ADHD score as simple effect.

Model*_Accuracy_ADHD_* ∼ Group * Temporal Variability + ADHD_score + (1|Participant)
Model*_RT_ADHD_* ∼ Group * Temporal Variability * Response Correctness + ADHD_score + (1|Participant)

We performed post-hoc tests using emmeans (package version 1.7.4.1.) with Tukey multiple comparison correction, to evaluate the difference in performance between Temporal Variability conditions. For this, we ran the model with Temporal Variability as a categorical factor with six levels (0, 25, 50, 75, 100, 150 ms Temporal *SD*). Stepwise models’ comparison was done using the likelihood ratio test, and Type II Wald χ^2^tests were used to assess the best model fit, and the significance of fixed effects (Bates et al., 2014; Luke, 2017).

## RESULTS

### Demographics

No significant differences were observed between the Dyslexic and Control groups regarding gender balance, age, years of post-baccalaureate study, and IQ as indicated by the Raven matrix score (Table 1). Although the participant with a score indicative of an ADHD disorder was excluded, dyslexic participants scored higher in the ADHD test than controls (Table 1). Participants with dyslexia demonstrated significant differences compared to the control group across all dyslexia screening tests (Table 1).

**Table 1:** Demographic characteristics and scores of the dyslexia screening tests in the control and dyslexic groups.

|  |  | Dyslexics | Controls | Statistics |
| --- | --- | --- | --- | --- |
| Demographics | Gender | 18 W/ 8 M | 19 W/ 7 M | $\chi^2(1) = 0, p > .99$ |
| | Age | $24.5 \text{ y} \pm 6.4 \text{ SD}$ | $24.2 \text{ y} \pm 4.5 \text{ SD}$ | $t(50) = -0.20, p = .84$ |
| | Years post-bac | $2.4 \text{ y} \pm 1.5 \text{ SD}$ | $3.2 \text{ y} \pm 1.6 \text{ SD}$ | $t(50) = 1.77, p = .08$ |
| | Non-verbal IQ | $112.2 \pm 11.0 \text{ SD}$ | $115.6 \pm 5.8 \text{ SD}$ | $t(50) = 1.42, p = .16$ |
| | ADHD score | $2.8 \pm 1.2 \text{ SD}$ | $1.5 \pm 1.3 \text{ SD}$ | $t(50) = -3.67, p < .001$ |
| Alouette | Corrected reading time | $146.5 \text{ s} \pm 37.0 \text{ SD}$ | $102.6 \text{ s} \pm 16.1 \text{ SD}$ | $t(50) = -5.49, p < .001$ |
| Initial phoneme deletion test | time | $48.1 \text{ s} \pm 14.7 \text{ SD}$ | $35.3 \text{ s} \pm 9.4 \text{ SD}$ | $t(50) = -3.25, p = .002$ |
| | score | $6.8 \pm 2.4 \text{ SD}$ | $8.69 \pm 1.5 \text{ SD}$ | $t(50) = 3.01, p = .004$ |
| Spoonerism test | time | $159.0 \text{ s} \pm 56.6 \text{ SD}$ | $87.3 \text{ s} \pm 27.1 \text{ SD}$ | $t(50) = -5.61, p < .001$ |
| | score | $15.8 \pm 4.6 \text{ SD}$ | $19.1 \pm 0.7 \text{ SD}$ | $t(50) = -3.73, p < .001$ |
| Dictation errors | Regular words | $2.7 \pm 1.8 \text{ SD}$ | $0.8, \pm 1.0 \text{ SD}$ | $t(50) = -4.92, p < .001$ |
| | Irregular words | $5.6 \pm 1.8 \text{ SD}$ | $2.2 \pm 2.0 \text{ SD}$ | $t(50) = -6.45, p < .001$ |
| | Pseudowords | $1.6 \pm 1.2 \text{ SD}$ | $1.0 \pm 0.7 \text{ SD}$ | $t(50) = -2.13, p = .04$ |

### Auditory discrimination task

Auditory discrimination thresholds were not statistically different between the two groups: participants staircases’ SNR thresholds were comparable (leading to an average target pitch discrimination of 80% in temporally irregular sound sequences) (*t*(50) = 0.78, *p* = .44, dyslexics: *M* = −14.1 dB ± 1.0 *SD*; controls: *M* = −13.9 dB ± 1.3 *SD*). Yet, dyslexic participants overall discriminated more poorly the deviant tones in the sequences than did the controls. Dyslexics had less correct responses (74.9 % ± 11.0 *SD*) than did controls (82.6 % ± 7.8 SD) (Fig. 2A-B, effect of the factor Group: χ^2^(1) = 6.57, *p* = .01), i.e. an average difference of 7.7 % between the groups. The variability of the context did not significantly affect response accuracy (Fig.2C, no effect of the factor Temporal Variability: χ^2^(1) = 2.00, *p* = .16), and no Group × Temporal Variability interaction (χ^2^(1) = 0.41, *p* = .52), suggesting that the accuracy of dyslexics and controls did not differ as a function of the temporal variability of the stimuli.

**Figure 2:**
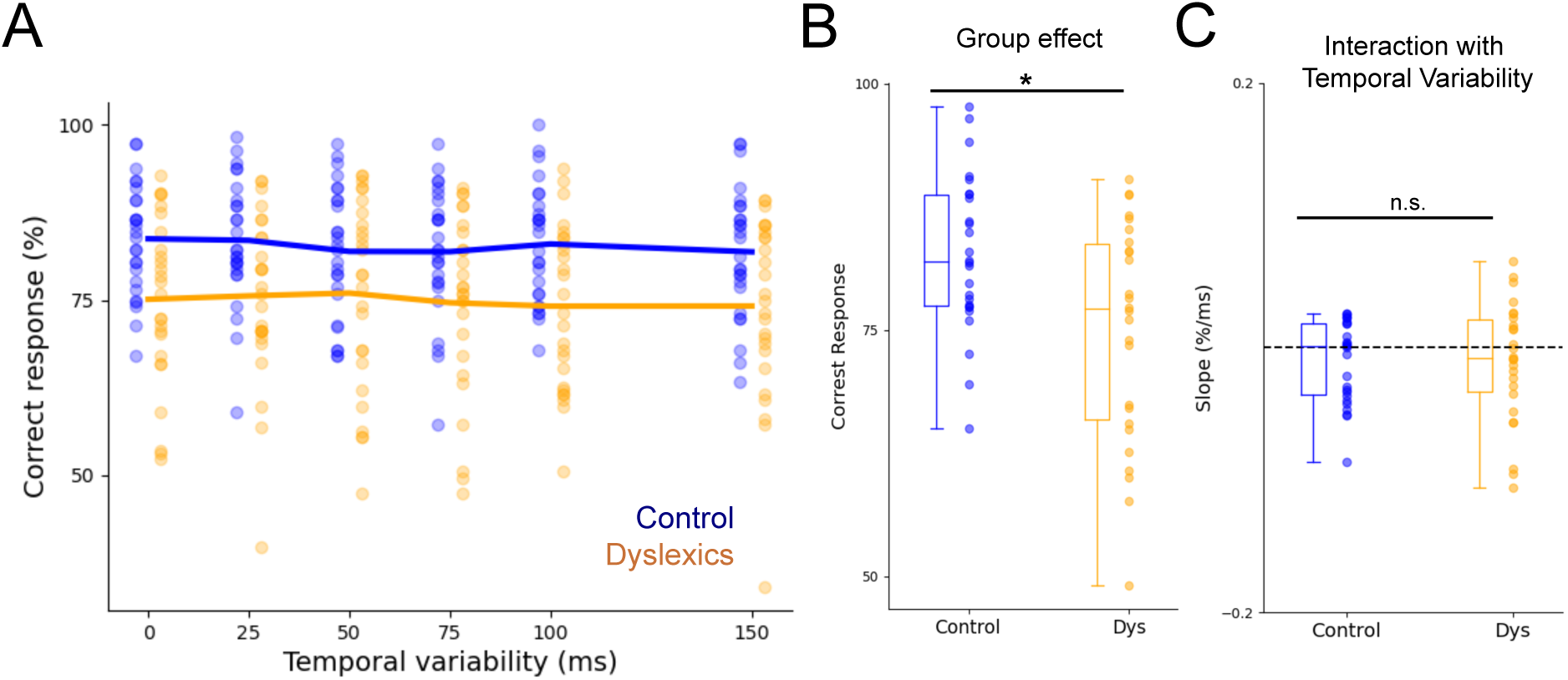
Auditory discrimination is affected by dyslexia. (A) Percentages of correct responses are represented as a function of the Temporal Variability (SD of SOAs) in the auditory sequences. Dots represent individual data and lines denote the mean across participants. (B) Control participants overall performed better than dyslexic participants. (C) Boxplots representing the coefficients of regressions of the percentage of correct response as a function of temporal variability for control and dyslexic participants. Auditory discrimination was not significantly affected by the temporal variability of the sequence in both groups.

RTs did not significantly differ between the groups (Fig. 3A-B, dyslexics: 670.5 ms ± 123.1 *SD*, controls: 635.0 ms ± 117.7 *SD*, χ^2^(1) = 0.72, *p* = .40). Participants overall responded faster when they gave a correct response, as compared to an incorrect response (χ^2^(1) = 199.97, *p* < .001), but this effect was dependent on the group (Response Correctness × Group: χ^2^(1) = 24.94, *p* < .001). The significant interaction suggests that the difference in RTs between correct and incorrect responses was smaller in the dyslexic group (- 35.7 ms ± 56.2 *SD*) than in the control group (- 71.0 ms ± 81.2 *SD*) (Fig. 3C).

**Figure 3:**
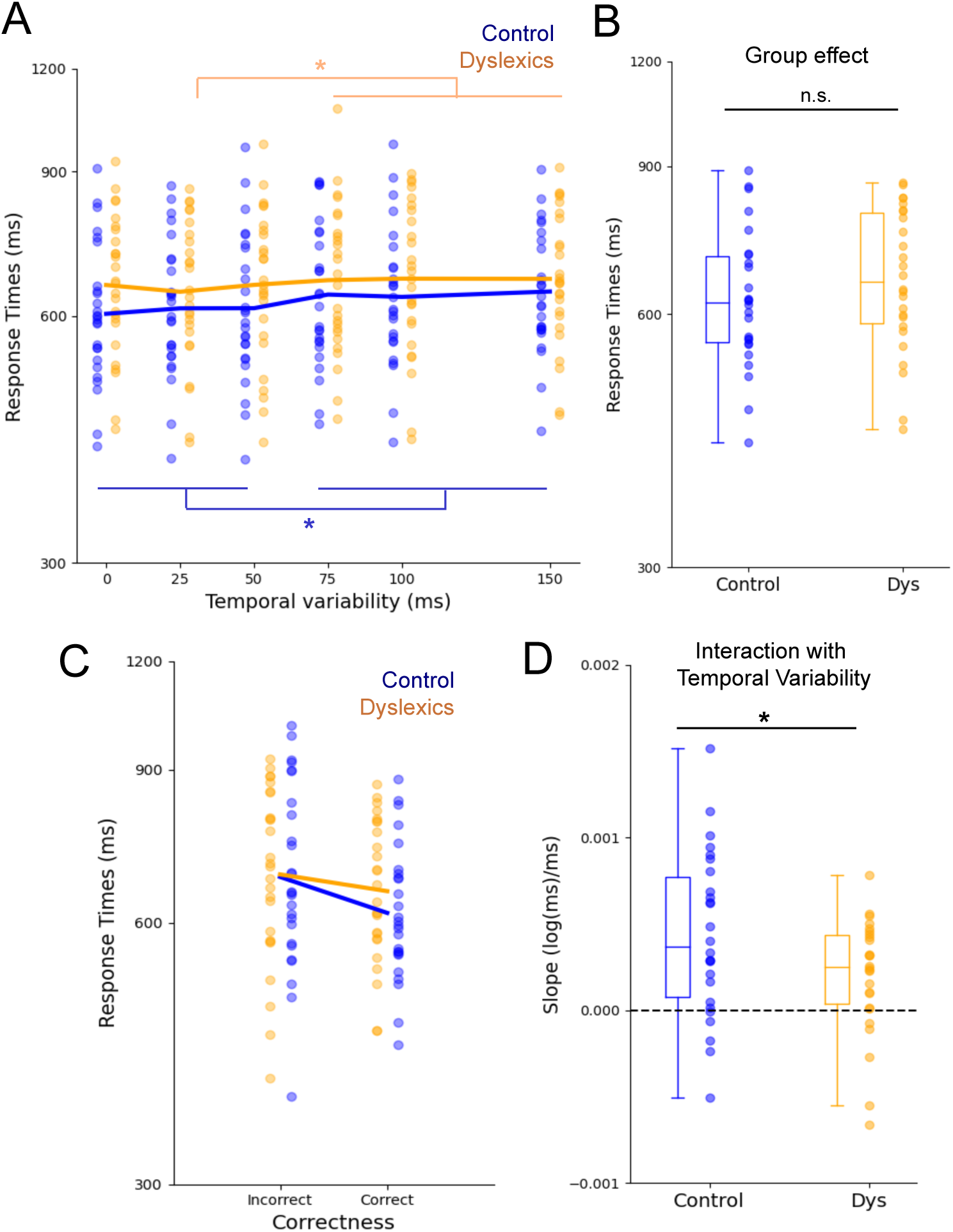
Variations in target’s response times (RT) as a function of temporal variability in control and dyslexic participants. (A) Log RTs as a function of the temporal variability of the auditory sequences. (B) RTs did not significantly differ between groups. (C) Participants overall responded faster to give correct responses than incorrect response, but this effect was smaller in the dyslexic group than in the control group. (D) Boxplots representing the coefficients of regressions of the log-transformed RTs as a function of temporal variability for control and dyslexic participants. RTs were more strongly affected by the temporal variability of the context (more positive RT slopes) in control participants than in dyslexic participants. A RT slope above 0 means that RTs become slower as the temporal variability of the context increases.

Importantly, the variability of the temporal context influenced RTs. Participants responded faster when the targets were embedded in more temporally regular sequences (effect of Temporal Variability: χ^2^(1) = 90.94, *p* < .001), and a significant Group × Temporal Variability interaction was observed (χ^2^(1) = 11.26, *p* < .001): the impact of temporal variability on RTs was lower in the dyslexia group than in the control group (Fig. 3D). Post-hoc tests showed that in controls, RTs were significantly faster if the sequence had low temporal variability (<75 ms *SD*) than higher temporal variability (≥75 ms *SD*) (0 ms / 75 ms *SD*: *p* < .001; 0 ms / 100 ms *SD*: *p <* .001; 0 ms / 150 ms *SD*: *p <* .001; 25 ms / 75 ms *SD*: *p* < .001; 25 ms / 100 ms *SD*: *p* = .003; 25 ms / 150 ms *SD*: *p <* .001; 50 ms / 75 ms *SD*: *p <* .001; 50 ms / 100 ms *SD*: *p =* .007; 50 ms / 150 ms *SD*: *p <* .001). These patterns of results were similar to previously reported findings (Bonnet et al., 2024). In dyslexic participants, RTs were only faster in the 25 ms *SD* condition compared to ≥75 ms *SD* conditions (25 ms / 75 ms *SD*: *p =* .03; 25 ms / 100 ms *SD*: *p =* .003; 25 ms / 150 ms SD: *p =* .001). Albeit with overall smaller slopes, the dyslexic group was also significantly influenced by the temporal context. Analyzing each group separately in two distinct GLMMs showed a significant effect of Temporal Variability in each group, (dyslexic: *χ^2^*(1) = 15.91; *p* < .001; control: χ2 (1) = 101.07; *p* < .001).

No significant Temporal Variability × Correctness interaction was observed (Correctness × Temporal Variability: *χ^2^*(1) = 0.46, *p* = .50; interaction Correctness × Temporal Variability × Group: *χ^2^*(1) = 0.54, *p* = .46), indicating that the influence of temporal variability on RTs was not significantly different between correct and incorrect responses. Additionally, although ADHD scores were different between our two groups, adding the ADHD score as a fixed simple effect in our statistical models for the discrimination task did not significantly improve the model fit for both response accuracy (AIC model_Accuracy_ = 30995, AIC model_Accuracy_ADHD_ = 30997, χ^2^(1) = 0.08; *p* = .77) and RTs (AIC model_Accuracy_ = −16582, AIC model_Accuracy_ADHD_ = 16581, χ^2^(1) = 0.12; *p* = .72).

### Subjective perception of regularity and participants’ performance estimations

The subjective perception of temporal regularity was influenced by the Temporal Variability (χ^2^(1) = 270.66, *p* < .001), with more regular sequences rated as more “regular” than irregular ones (Fig. 4A). No overall between-group difference in rating was found (χ^2^(1) = 0.51, *p* = .47) (Fig. 4B), but the Group × Temporal Variability interaction was significant (χ^2^(1) = 12.97, *p* < .001): compared with the control group, dyslexic participants overestimated the temporal regularity of irregular contexts and underestimated that of regular ones (Fig. 4A-C).

**Figure 4:**
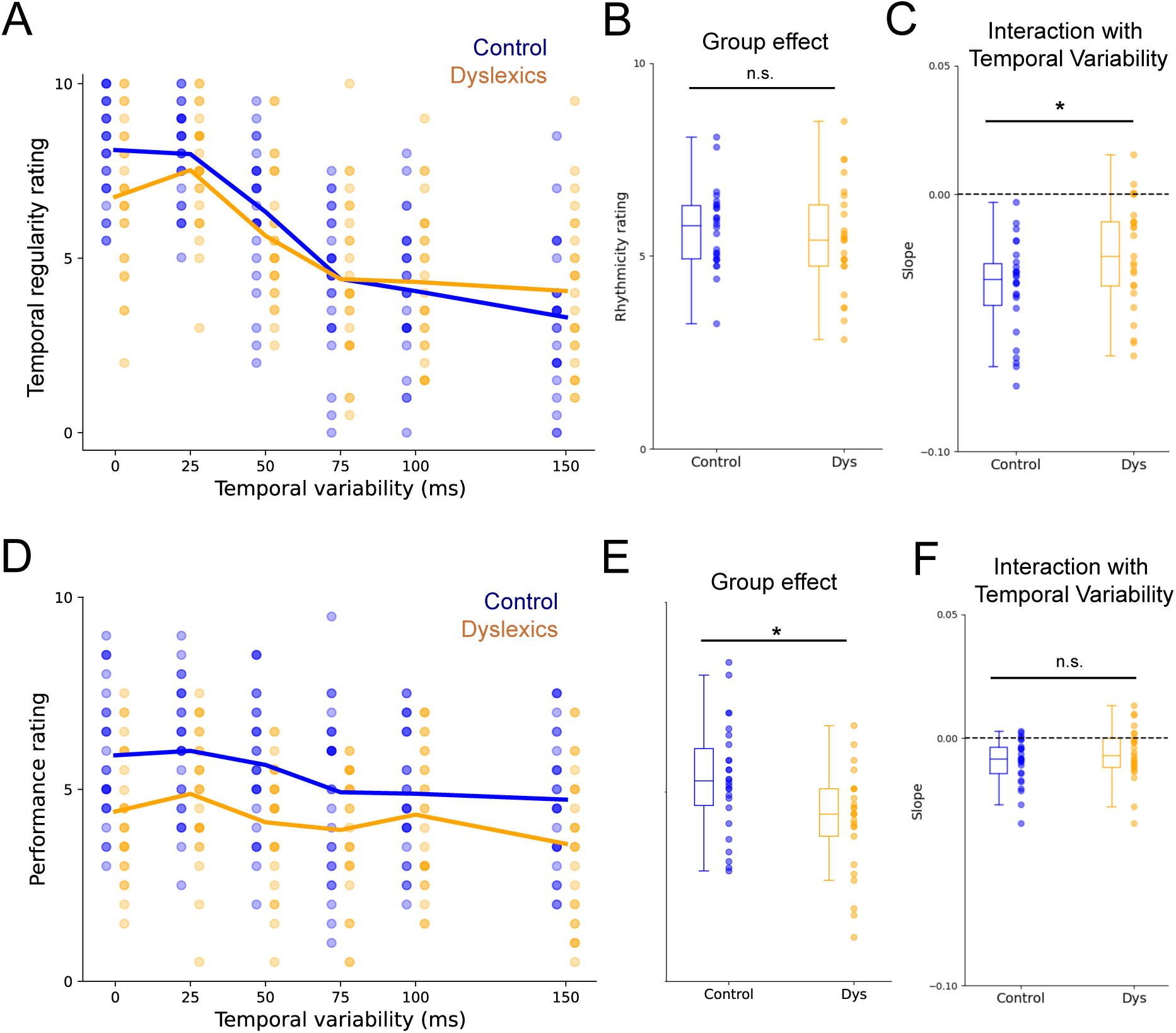
Subjective ratings of temporal regularity and performance were influenced by the temporal regularity of sequences in dyslexia. (A) Rating of temporal regularity of sound sequences, dots represent individual data and lines denote the mean across participants. Both groups perceived the more temporally regular sequences as more “regular”. (B) Overall judgments of temporal regularity did not change between groups. However, (C) compared with the control group, dyslexic participants overestimated the temporal regularity of irregular contexts and underestimated that of regular contexts. (D) Subjective performance ratings. (E) Dyslexic participants reported lower overall performance ratings throughout the task. (F) Both groups perceived their auditory discrimination abilities as lower when engaging with sequences exhibiting the greatest temporal variability; however, no significant Group × Temporal Variability interaction was observed.

Participants’ subjective ratings about their own performance appeared to be affected by the temporal variability of the context (Fig. 4D, χ^2^(1) = 37.15, *p* < .001). Specifically, participants perceived themselves to be more accurate in discriminating sounds within more regular contexts, consistent with the RT results indicating faster discrimination, but not with their accuracy (no effect of temporal variability on proportion of correct responses; Fig. 2A,C). A difference was found between the groups (*χ^2^*(1) = 8.15, *p* < .001), but without significant Group × Temporal Variability interaction (χ^2^(1) = 1.55, *p* = .21). This result suggests that dyslexic participants had overall lower confidence in their performance throughout the task.

## DISCUSSION

Our present study aimed to assess the influence of probabilistic temporal regularities in auditory sequences on temporal predictions and on sound pitch processing. Results provide converging evidence with our previous research on typical adults (Bonnet et al., 2024), revealing that variability in temporal context influences auditory discrimination performance (here on the pitch dimension). Participants showed slower responses to target sounds when engaged with sound sequences that featured greater temporal variations. Importantly, our data revealed a significant disparity between the two groups of participants. Dyslexic individuals demonstrated lower accuracy in pitch discrimination of the target sounds, across all presented temporal sequences, when compared to control participants. Furthermore, the influence of temporal variability on response speed exhibited a distinct pattern for each group. Control participants demonstrated a greater sensitivity to temporal variability, with their response times increasing correspondingly as the temporal variability of the sequence grew, whereas individuals with dyslexia showed a comparatively lesser effect. These findings suggest that dyslexic adults may encounter greater challenges in processing sequential auditory information and, unlike control participants, derive less benefit from temporal context regularity.

### Inefficient Temporal Processing and Prediction in Dyslexia

Our findings indicate a significant overall difficulty among participants with dyslexia in distinguishing deviant sounds within continuous sound sequences. This result pattern was observed despite participants of both groups displaying comparable perceptual auditory thresholds in the staircase procedure. These findings are consistent with recent research indicating that individuals with dyslexia may face a generalized challenge in auditory statistical learning, such as when processing the transitional probabilities between tones presented in sequences (Gabay et al., 2015, 2023; Ozernov-Palchik et al., 2023; Ringer et al., 2024). These results fit with the anchoring-deficit theory (Ahissar et al., 2000, 2006; Daikhin et al., 2017), which proposes that, in typical neurological functioning, repeated exposure to a specific auditory reference enhances the ability to discriminate an unexpected deviant sound. However, individuals with dyslexia seem to possess a compromised mechanism in building sound categories (Serniclaes et al., 2004), resulting in reduced advantages from such repetition. This may account for the overall decrease in auditory discrimination performance observed in our study in participants with dyslexia: control participants likely benefitted from the repetition of standard tones when identifying deviants, while dyslexic participants did not experience the same amount of advantage. These patterns of results could also reflect impaired memory trace of standard and deviant tones, and be linked to non-verbal short-term memory deficits observed in dyslexia (Trecy et al., 2013).

Our data further revealed that the temporal variability of sound sequences differentially influenced auditory perception in the two groups of participants: the temporal variability of the auditory sequences affected response speed in control groups, with faster responses in more temporally predictable sequences. This effect was less strong in the dyslexic group, suggesting that their response speed was less influenced by the temporal variability of the sequences. These results can be interpreted within the context of temporal impairment frameworks related to dyslexia (Goswami, 2011, 2018; Tallal et al., 1993), but here showing a specific effect on temporal predictability, which was not task-relevant (Taha, et al., 2025). Dyslexic participants may not utilize temporal prediction cues as effectively as their control counterparts, and therefore they may respond relatively independently of the temporal context. It might be argued, in line with the sluggish attentional shifting theory (Hari & Renvall, 2001; Lallier et al., 2010), that dyslexic individuals’ automatic attention system struggles to disengage promptly from one stimulus to move on to the next encounter. This effect has been observed for rapid stimulus sequences (<200 ms SOA) (Hari & Renvall, 2001; Lallier et al., 2010). However, here, we presented slower tempos (500 ms) to participants. Our results could therefore suggest that not only the speed of sequences, but also their temporal variability, could affect temporal attention mechanisms. We further argue that these findings are specifically related to temporal attention mechanisms rather than broader domain-general deficits in attention associated with dyslexia. Although significant differences in ADHD scores were observed among the groups, including ADHD scores as a fixed effect did not enhance the models’ ability to account for the behavioral data.

### Implications for Neural Oscillatory Frameworks in the Context of Developmental Dyslexia

Multiple models indicate that dyslexia may stem from deficits in neural synchronization with the slow timing dynamics of speech (Goswami, 2011, 2018; Lallier et al., 2017; Leong, et al., 2014). Brain oscillations tend to align with the rhythmic amplitude modulations of speech. This speech-brain synchronization is believed to reflect mechanisms of auditory attention regulation. The synchronization of low-frequency oscillations facilitates attentional shifts between prominent prosodic units, which in turn guide subsequent phoneme sampling that is essential in developing phonological skills necessary for reading acquisition (Ziegler, Perry, & Zorzi, 2014). Neural mechanisms responsible for tracking rhythmic patterns must adapt to temporal variability to facilitate optimal processing of upcoming events (Zoefel & Kösem, 2024). This is relevant for human speech, which frequently exhibits variable rates and significant temporal fluctuations (Tilsen & Arvaniti, 2013; Varnet et al., 2017). Prior neuroimaging research has demonstrated that neural dynamics involved in auditory rhythm tracking adapt based on the statistical properties of previous auditory input (Bouwer et al., 2023; van Bree et al., 2021; Kösem et al., 2018). Our data can be integrated in this view by showing that the perception of auditory rhythms is context-dependent and influenced by the temporal statistics of prior stimuli. The findings further suggest that individuals with dyslexia and neurotypical controls may differ in their neural capacities to track temporal patterns amid variability. Notably, Fiveash and colleagues (2020) recently reported reduced neural synchronization to complex, irregular rhythms in adults with dyslexia compared to controls. These findings align with our results, indicating that control participants are better able to process subtle variations in temporal regularities in continuous sounds even when presented in jittered sequences, whereas individuals with dyslexia may exhibit difficulties in this regard.

### Subjective Perception of Temporal Regularity and Confidence Decline in Dyslexia

Compared with the control group, the dyslexic participants overestimated the temporal regularity of irregular contexts and underestimated that of regular ones. These findings align with previous reports suggesting impaired judgments in explicit temporal tasks in dyslexic adults and children (Bégel et al., 2022; Canette et al., 2020; Leong & Goswami, 2014). Moreover, we also reported that subjective perception of auditory discrimination performance was impacted in dyslexia, showing that dyslexics participants have overall lower confidence in their answers to the task, again in line with preceding findings (Canette et al., 2020). This finding may indicate a domain-specific decrease in confidence related to temporal tasks among individuals with dyslexia. Conversely, the lower confidence ratings observed in participants with dyslexia could also stem from a broader, domain-general negative self-perception of their learning abilities, as has been documented in academic contexts (Dykes, 2019).

### Effect of Music (Rhythmic) Training and Rhythm Exposure on Auditory Perception

Our findings do not fully align with those of Bonnet et al. (2024). Whereas the preceding study reported effects on both speed and accuracy of responses, the present data showed an effect of temporal variability only for response times, including among control participants. One potential explanation for this effect could be the role of musical practice. In our previous study we did not control for musical practice therefore potentially including musicians in our experiment, while in the present study we purposely included participants who had no musical experience, which may suggest that individuals without musical training may exhibit diminished temporal prediction abilities. Musical practice is known to influence the perception of rhythm (Tallal & Gaab, 2006), and neural tracking to rhythms (Musacchia et al., 2008). Musical practice may also influence speech perception, particularly in dyslexics (Flaugnacco et al., 2015; Habib et al., 2016). It is therefore possible that musical practice, and training aimed at enhancing rhythm processing, may offer a valuable method for addressing language processing difficulties associated with dyslexia. However, it is important to acknowledge that while musical training may improve speech processing abilities, it does not eliminate the risk of reading disorders. As a result, individuals who have undergone musical training may still encounter ongoing challenges associated with dyslexia (Bishop-Liebler et al., 2014; Couvignou, et al., 2024; Zuk et al., 2017).

Rhythmic priming studies, which involve brief exposure to musical rhythms prior to speech presentation, have demonstrated an influence on speech perception across various populations of adults and children, including individuals with and without language disorders (Bedoin et al., 2016; Camici et al., 2025; Canette et al., 2020; Fiveash et al., 2020b, 2023; Przybylski et al., 2013). Notably, rhythmic priming appears to be most effective when the rhythmic structure in the musical stimulus is sufficiently long and salient (Fiveash et al., 2020b). Our findings support the notion that, for rhythmic priming to be effective, the rhythmic pattern must be salient, as individuals with dyslexia may struggle to perceive the underlying structure of complex rhythms and temporal contexts with enhanced temporal variability.

## CONCLUSION

Our study provides converging evidence for altered temporal prediction in dyslexia, for both implicit temporal processing that influences pitch discrimination performances, and explicit temporal regularity judgments. Our findings demonstrate that individuals with dyslexia show reduced sensitivity to temporal regularities and irregularities in auditory sequences. These deficits impair dyslexic individuals’ ability to accurately perceive and process sounds in complex rhythmic sequences, potentially contributing to difficulties in continuous speech processing. The findings emphasize the significance of temporal context in auditory perception and provide valuable insights into the underlying mechanisms that may influence language-related skills in individuals with dyslexia.

## ACKNOWLEDGMENTS

This work was supported by an ANR grant (ANR-21-CE37-0003) to A.K. BT was supported by an ANER grant of the Region Bourgogne-Franche Comté (MusiC). We would like to thank Lauriane Perpere for her help with data acquisition.

